# High temporal resolution reveals simultaneous plasma membrane recruitment of the TPLATE complex subunits

**DOI:** 10.1101/2020.02.13.948109

**Authors:** Jie Wang, Evelien Mylle, Alexander Johnson, Nienke Besbrugge, Geert De Jaeger, Jiří Friml, Roman Pleskot, Daniel Van Damme

## Abstract

The TPLATE complex (TPC) is a key endocytic adaptor protein complex in plants. TPC contains six evolutionary conserved subunits and two plant specific subunits, AtEH1/Pan1 and AtEH2/Pan1, which are not associated with the hexameric subcomplex in the cytoplasm. To investigate the dynamic assembly of the octameric TPC at the plasma membrane (PM), we performed state-of-the-art dual-color live cell imaging at physiological and a lowered temperature. Our data show that lowering the temperature slows down endocytosis and thereby enhances the temporal resolution of the differential recruitment of endocytic components. Under both normal and lowered temperature conditions, the core TPC subunit TPLATE, and the AtEH/Pan1 proteins, exhibited simultaneous recruitment at the PM. These results, together with our co-localization analysis of different TPC subunits, allow us to conclude that in plant cells, TPC is not recruited to the PM sequentially but as an octameric complex.

**One sentence summary:** Lowering the temperature increases spatiotemporal resolution of protein recruitment at the plasma membrane.

## Introduction

Clathrin-mediated endocytosis (CME) is the best-studied and predominant endocytic pathway in eukaryotes to internalize plasma membrane (PM) proteins and extracellular materials, commonly termed “cargo”, into cells (Bitsikas et al., 2014; Dhonukshe et al., 2007; Kitakura et al., 2011; Robert et al., 2010). The formation of clathrin-coated vesicles (CCVs) requires several highly coordinated stages: initiation, cargo selection, coat assembly, scission and vesicle uncoating (McMahon and Boucrot, 2011). Though the initiation of CME at the PM remains poorly understood in plants, similarly to other systems, adaptor proteins are presumed to recognize cargo proteins via cargo-recognition motifs and subsequently recruit clathrin triskelia through clathrin-binding motifs (Zhang et al., 2015). As clathrin does not bind directly to the PM nor to the cargo proteins, adaptor proteins thus play an essential role to link the PM and the clathrin cage (McMahon and Boucrot, 2011).

Two early-arriving adaptor protein complexes function at the PM in plants; the heterotetrameric Adaptor Protein-2 complex (AP-2) and the octameric TPLATE complex (TPC). AP-2 comprises of two large (AP2A and AP2B or α and β), one medium (AP2M or μ) and one small subunit (AP2S or σ) (Di Rubbo et al., 2013). TPC contains eight components; evolutionary conserved TPLATE, TML, TASH3, LOLITA, TWD40-1, TWD40-2, and plant-specific AtEH1/Pan1 and AtEH2/Pan1 subunits (Gadeyne et al., 2014; Hirst et al., 2014). AP-2 function is important for somatic plant development as single *Arabidopsis ap-2* mutants display developmental defects, yet still result in viable plants (Di Rubbo et al., 2013; Fan et al., 2013; Kim et al., 2013; Yamaoka et al., 2013). The TPC is essential for both pollen and somatic plant development as knockout, or knockdown, of single subunits of TPC in *Arabidopsis* leads to pollen or seedling lethality respectively (Gadeyne et al., 2014; Van Damme et al., 2006).

TPC is an evolutionary ancient protein complex that has been so far experimentally characterized only in plants and in an amoeba, *Dictyostelium discoideum* (Gadeyne et al., 2014; Hirst et al., 2014). TPC in plants was identified as an octameric complex by tandem affinity purification (TAP) experiments (Gadeyne et al., 2014). Recently, we also identified an important role for the AtEH/Pan1 proteins in actin-regulated autophagy and in recruiting several components of the endocytic machinery to the AtEH/Pan1 positive autophagosomes (Gadeyne et al., 2014; Wang et al., 2019). TPC in *Dictyostelium* (described as TSET) was however identified as a hexameric complex lacking the AtEH/Pan1 homologs (Hirst et al., 2014). Although Amoebozoa contain homologuous proteins to AtEH/Pan1, these resemble more closely Ede1 than Pan1 (Gadeyne et al., 2014; Wang et al., 2019).

Truncation of the TML subunit of TPC forces the complex into the cytoplasm and this correlates with the dissociation of AtEH/Pan1 proteins from the complex (Gadeyne et al., 2014). It therefore appears that TPC composition depends on its localization and that the two AtEH/Pan1 subunits might be peripherally associated with a hexameric subcomplex of TPC, which would resemble to TSET in *Dictyostelium*. To investigate whether there is a differential order of recruitment between both AtEH/Pan1 proteins and the remaining hexameric subcomplex, we performed dual color time-lapse microscopy of CME in etiolated *Arabidopsis* epidermal hypocotyl cells. We also took advantage of lowering the temperature of our samples to slow down CME and thereby increase the spatiotemporal resolution. Altogether, our data strongly suggest that TPC is recruited as the octameric complex to the PM, where it functions as the early adaptor complex for plant CME.

## Results

### Lowering the experimental temperature reduces CME kinetics

TPC is proposed to serve as an early adaptor complex (Gadeyne et al., 2014), however temporal resolution however remains the biggest challenge to monitor the dynamic recruitment of different endocytic protein players (Bashline et al., 2015; Fan et al., 2013; Fujimoto et al., 2010; Gadeyne et al., 2014). As lowering the temperature generally slows down dynamics of cellular processes (Das et al., 1966), we hypothesized that by lowering the temperature, we could also slow down dynamics of endocytic events and therefore enhance the spatiotemporal visualization of the differential recruitment of the endocytic players at the PM.

To time how intracellular dynamics respond to decreasing the temperature in plant cells, we firstly imaged *Arabidopsis* plants expressing the microtubule (MT) binding protein EB1a-GFP (Van Damme et al., 2004). Hypocotyl cells of *Arabidopsis* seedlings expressing EB1a-GFP were imaged at 20 °C for 5 min, then the temperature was lowered to 10 °C with the aid of the CherryTemp Heater Cooler system and the effect on microtubule polymerization rate was assessed. Lowering the temperature had an immediate effect on the MT polymerization speed, measured as EB1a labeled tracks in time-projection images (Fig. S1 A). Visualization and quantification of the MT growth dynamics further confirmed that the temperature shift was transmitted to the seedling almost instantaneously (Fig. S1).

To examine the capacity of lowering the temperature to slow down endocytosis, etiolated hypocotyl cells expressing endocytic markers CLC2-mKO and TPLATE-GFP the temperature was shifted during image acquisition and no obvious defect in the recruitment was observed. Density analysis of early-arriving (TPLATE and TML) as well as late arriving DRP1A furthermore showed that lowering the temperature did not visually affect the PM recruitment of these endocytic markers while it prolonged their lifetime at the PM instantly (Figure 1 and Figure 2A).

**Figure 1.**
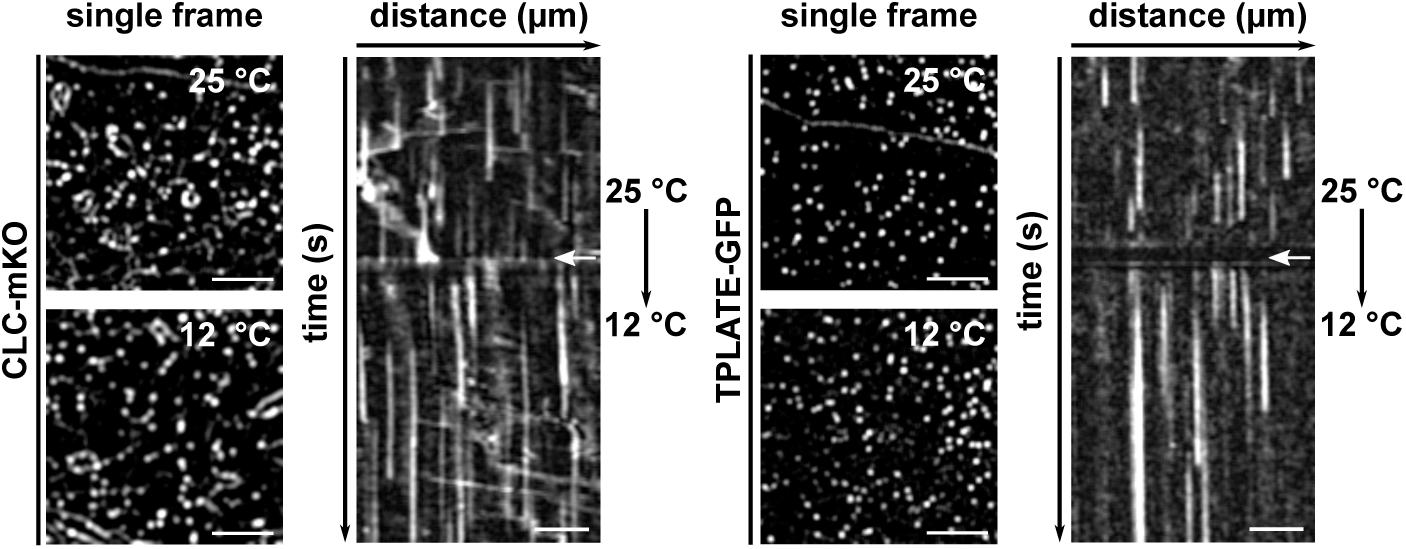
Endocytic dynamics alter immediately upon lowering the temperature. Spinning disc images and representative kymographs showing endocytic foci and lifetimes of CLC2 (left) or TPLATE (right) in Arabidopsis etiolated hypocotyl epidermal cells at different temperatures. Cells were imaged at 25 °C for 2 minutes and then imaged at 12 °C for an additional 3 minutes. Images were acquired with a 1-second interval. White arrows on the kymographs indicate the position of the temperature shift. Scale bars of spinning disc images, 5 µm. Scale bars of kymographs, 30 µm. Time, 300s.

**Figure 2.**
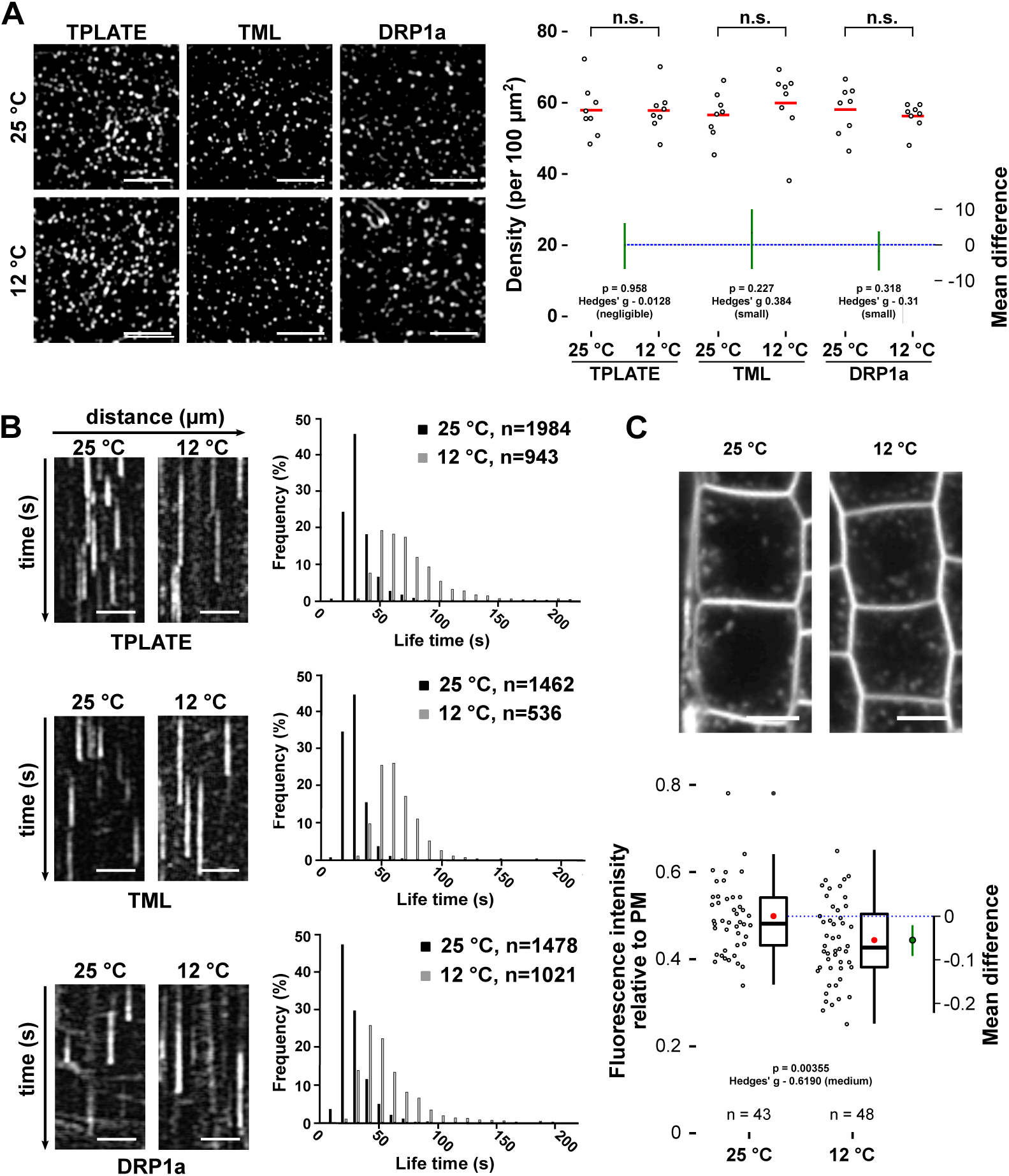
Lowering the temperature slows down CME. (A) Spinning disk images and density quantification of TPLATE, TML and DRP1a-marked foci in epidermal hypocotyl cells at the permissive (25 degrees) and restrictive (12 degrees) temperature. The density of the foci is independent of the tested temperatures (n.s. not significant, Mann-Whitney U-test). The mean difference with the bootstrap 95% confidential interval (green circle and green line) is shown as a part of the plot. Hedges’ g value is a standardized effect size. For each transgenic line, 8 cells from 8 seedlings were imaged at both temperatures. Scale bars, 5 µm. (B) Representative kymographs and histograms showing the life-time distribution of TPLATE, TML and DRP1a positive foci at both temperatures. Scale bars, 25 µm. Time, 120 s. Number of events (n) is indicated. (C) Representative confocal images and Jitter and box plot quantification showing internalization of the FM4-64 dye (4μM, 30mins) in Col-0 root epidermal cells at both temperatures. The red circle represents the mean. The p-value was calculated by the Welch two sample T-test. The mean difference with the bootstrap 95% confidential interval (green circle and green line) is shown on the right side of the plot. Hedges’ g value is a standardized effect size. n represents the number of measurements (n = 43 cells for 25 °C and n = 48 cells for 12 °C) from 11 individual roots respectively. Scale bar, 10 µm.

To further evaluate how lowering the temperature affects the dynamic behavior of endocytic proteins, we carefully measured the dynamics of TPLATE, TML and DRP1a at different temperatures. Kymograph and histogram analysis of measured lifetimes confirmed that lowering the temperature from 25 to 10 °C correlated with a gradual increase in lifetime of endocytic proteins at the PM (Fig. S2). For TPLATE and TML, we observed a strong increase in average lifetime between 15 and 12 °C. Therefore, we selected 25 °C and 12 °C as the two temperatures for our future experiments.

Visualizing the individual lifetimes of the three markers at both temperatures in a histogram showed a clear shift of the distribution to longer lifetimes (Fig. 2B) and this correlated with a significant reduction in the internalization of the styryl dye FM4-64, which is used as a proxy for endocytic flux (Bashline et al., 2013; Dejonghe et al., 2016; Dejonghe et al., 2019; Fan et al., 2013; Gadeyne et al., 2014; Van Damme et al., 2011) (Fig. 2C). Furthermore, photo-bleaching experiments showed a dramatic reduction in the recovery of endocytic foci at PM at the low temperature, in agreement with reduced dynamics of the process (Fig. S3). These results together, show that lowering the temperature in etiolated seedlings using our experimental setup slows down CME efficiently and rapidly.

### Lowering the temperature enhances the temporal resolution of plasma membrane recruitment

Having established the effect of lowering the temperature on endocytic dynamics in *Arabidopsis* cells, we then examined whether we would be able to enhance the temporal difference between an early (TPLATE) player and a late one (DRP1A). We therefore generated a dual marker line of TPLATE-GFP (Van Damme et al., 2006) and DRP1a-mRFP (Mravec et al., 2011) and compared their temporal behavior at both temperatures. When imaged at 25 °C, the time-lapse images and kymographs showed multiple independent events where TPLATE-GFP foci clearly appeared earlier than DRP1a-mRFP at the PM, while they disappeared together (Fig. 3A, B and Fig. S4). Lowering the temperature prolonged the lifetime of both TPLATE-GFP and DRP1a-mRFP (Fig. 3A-B), whereas their departure remained synchronized. Overall, the temporal difference between their PM recruitment was dramatically enhanced (Fig. 3A, B and Fig. S4), which resulted in a very significant mean difference of the paired lifetimes and the large Hedge’s g value, which indicates the effect size (Fig. 3C).

**Figure 3.**
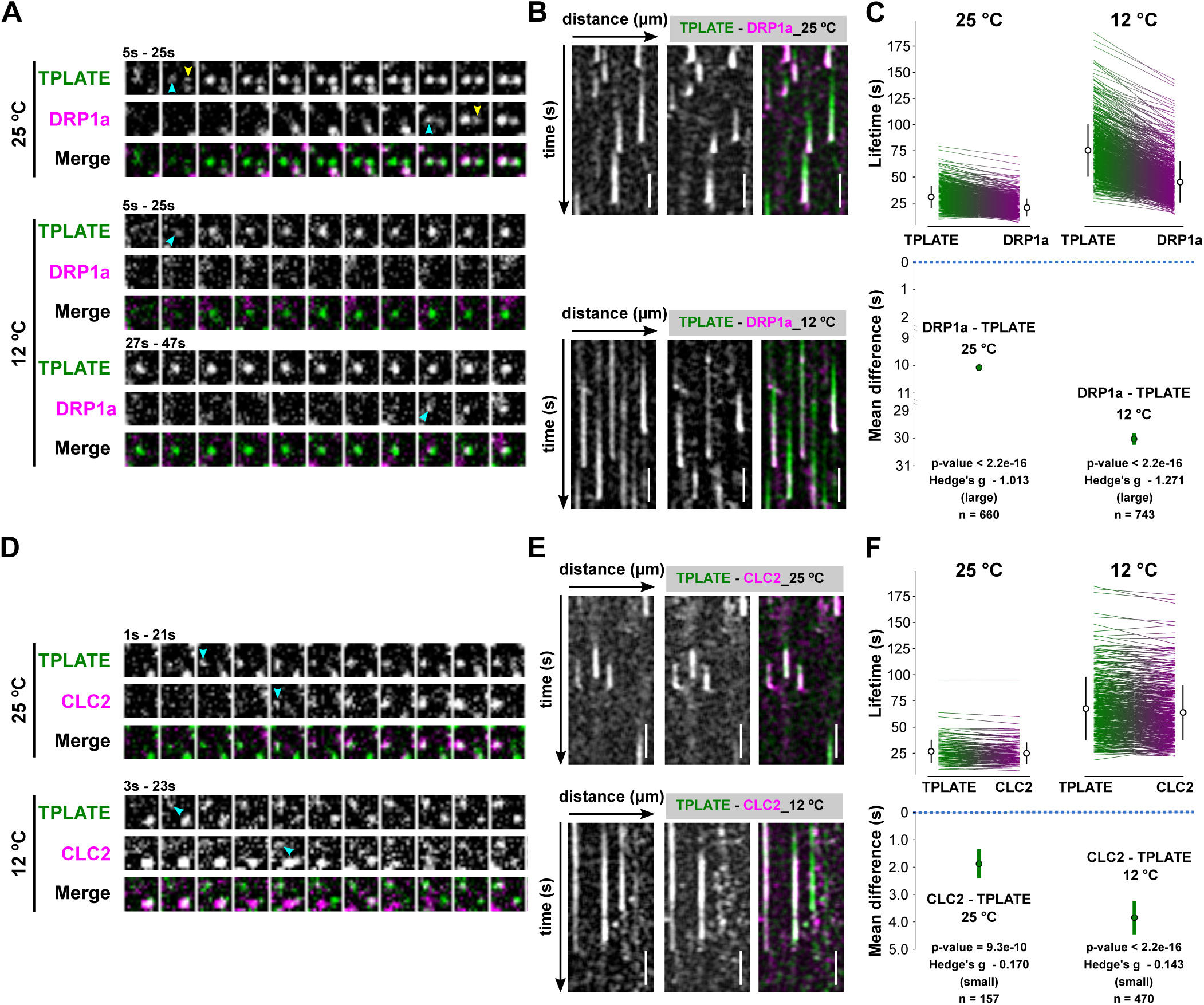
Lowering the temperature enhances the temporal resolution of plasma membrane recruitment. (A and D) Representative time series of dual-color spinning disk movies (2s/f) showing the sequential recruitment between TPLATE and DRP1a or CLC2 at the different temperatures. Arrowheads mark the appearance of TPLATE or DRP1a and CLC2 at PM. (B and E) Representative kymographs displaying the differential recruitment between TPLATE and DRP1a or CLC2 at different temperatures. Scale bar, 25 µm. Time, 120s. (C and F) Paired comparison of the lifetimes of particular protein pairs at the different temperatures. Each line represents an individual pair with the mean (white circle) ± SD (black line) on the sides of each plot for different combinations of constructs and temperatures. The p-value was calculated using the Wilcoxon Signed Rank test. The green circle in the bottom graph represents the paired mean difference with the bootstrap 95% confidential interval (green line). Hedges’ g value is a standardized effect size. n represents the number of events.

We performed a similar experiment comparing TPLATE and CLC2, which were previously shown only to have minor differences in their PM recruitment (Gadeyne et al., 2014; Narasimhan et al., 2020). Here, we also observed a temperature-enhancing effect of their differential recruitment with TPLATE being recruited before CLC2, yet less pronounced than the difference between TPLATE and DRP1a (Fig. 3D to F and Fig. S4).

Taken together, these results show that lowering the temperature slows down dynamics of endocytosis as well as enhances the spatiotemporal resolution of recruitment of the endocytic proteins at the PM.

### TPLATE is closely associated with the AtEH/Pan1 proteins at the plasma membrane

The absence of AtEH1/Pan1 and AtEH2/Pan1 as subunits of TSET (Hirst et al., 2014), together with the observation that inducing mislocalization of TPC to the cytoplasm leads to loss of those two subunits from the complex (Gadeyne et al., 2014) and the observation that AtEH/Pan1 proteins have a specific role in promoting autophagy (Wang et al., 2019), suggest that TPC could be in essence a hexameric complex, which temporarily gains two additional subunits. To reveal the spatiotemporal relationship among the TPC subunits, we visualized their dynamic behavior at the PM using multiple dual-color labeled lines. To evaluate the functional association of TPC subunits at PM, we crossed complemented mutant lines of core subunits (TPLATE, TML and TWD40-1) and generated double complemented homozygous mutant lines. We also combined complemented mutants of the core subunit TPLATE with AtEH1/Pan1 or AtEH2/Pan1 and generated the respective double complemented, double homozygous mutant lines. As a first indication for differential recruitment, we monitored the steady-state percentage of co-localization between the markers using a flattened projection of five consecutive frames. Comparing an early endocytosis marker such as TPLATE, with a late marker such as DRP1A revealed that the differential arrival between those makers leads to a substantial non-colocalizing fraction (roughly 40%, Fig. 4A).

**Figure 4.**
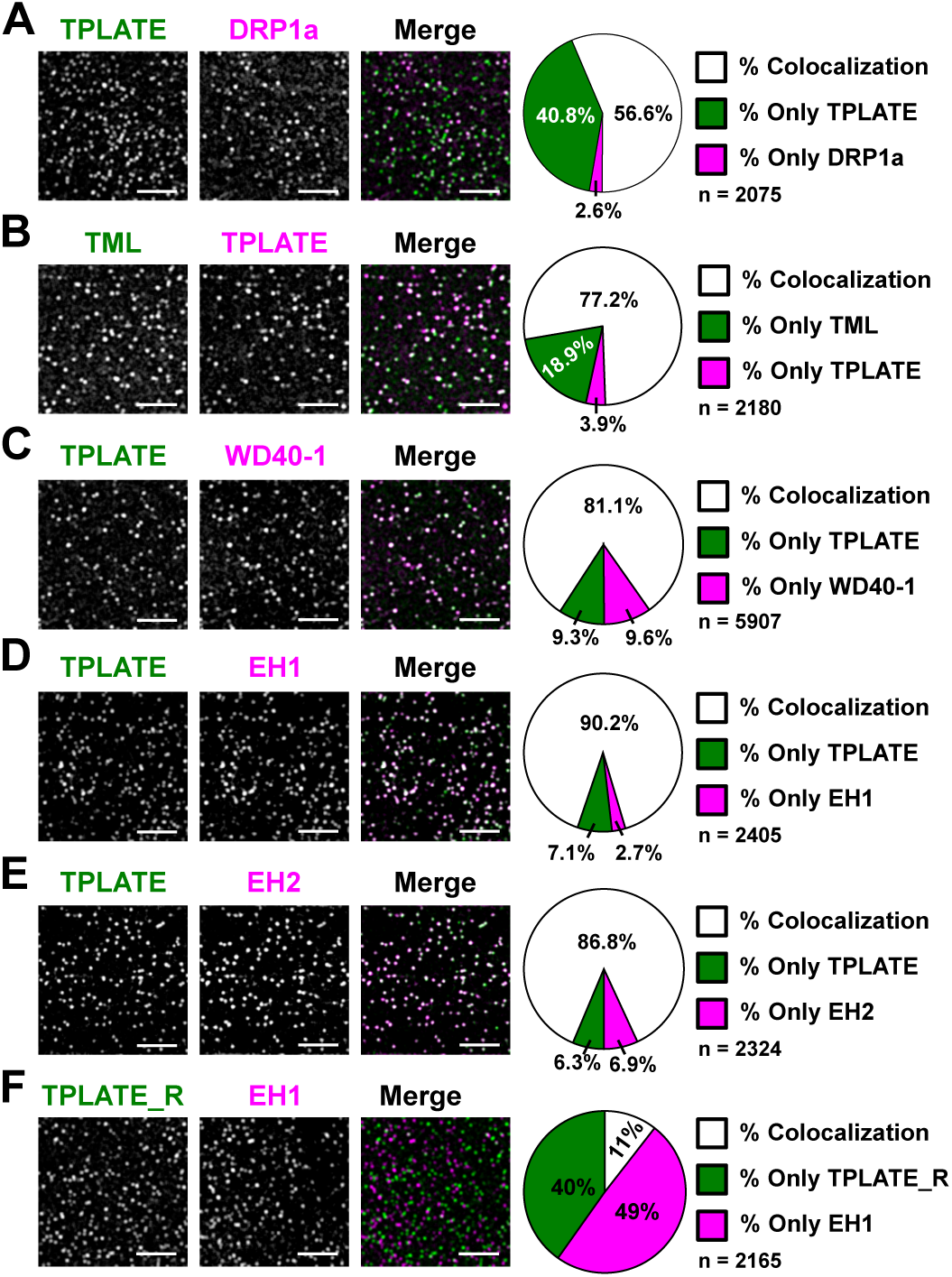
Colocalization analysis hints at a tight association between core and peripheral TPC subunits at the PM. Representative spinning disk dual-color images and corresponding pie charts displaying the percentage of colocalization of dual-labeled endocytic foci. TPLATE was compared to a late endocytic marker, DRP1a (A), to both TML (B) and TWD40-1 (C), and to both AtEHs/Pan1 (D, E). Rotated TPLATE (90° rotation; TPLATE_R) was also compared to AtEH1/Pan1 to control for random association of foci (F). Scale bars, 5 µm. Z-projections of five consecutive frames with average intensity were used for quantification. Quantification (%) of colocalization as calculated using the DIANA algorithm. The high percentage of overlap between TPLATE and the AtEH/Pan1 proteins suggests a tight connection between those proteins at the plasma membrane.

Combining TPLATE-TagRFP with TML-GFP resulted in around 77% of total foci where TML and TPLATE overlapped (Fig. 4B). We also tested functional association when combining TPLATE-GFP with a TWD40-1-mRuby3 complemented line (Fig. S5 and Table 1). Similarly, as for TML and TPLATE, combining TPLATE-GFP and TWD40-1-mRuby3 resulted in a very high percentage of colocalizing foci (Fig. 4C), confirming their intrinsic behavior as part of the same complex.

**Table 1.**
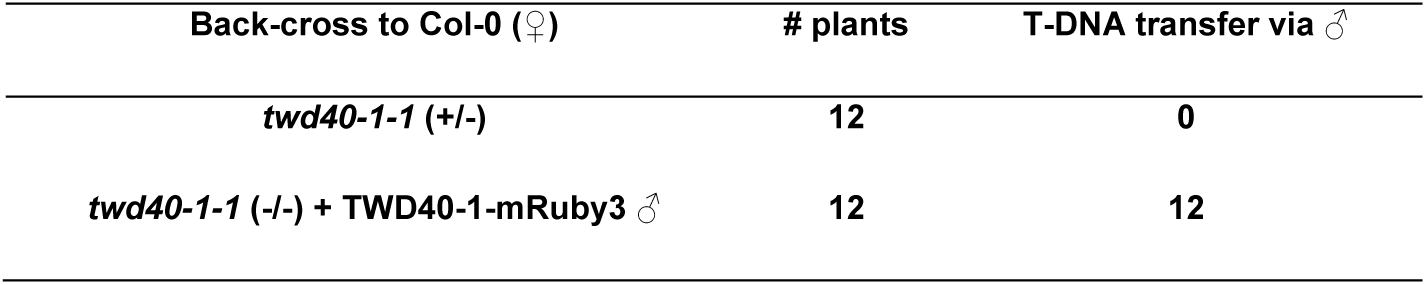
*twd40-1-1* plants expressing C-terminal fusions of TWD40-1 with mRuby3 allow transfer of the T-DNA via the male. The table shows the result of the analysis of back-cross experiments between Col-0 as female (♀ and the *twd40-1-1* heterozygous plants with or without expression of TWD40-1-mRuby3 as male (♂). The analysis clearly shows that the mutation blocks transfer via the male and that this block is lifted by the presence of the fluorescent fusion construct, indicating that it is functional.

In *Arabidopsis* seedlings, AtEH1/Pan1 and AtEH2/Pan1 have so far been shown functionally associated with TPLATE at autophagosomes and to be delivered to the vacuole after carbon starvation and Conc A treatment (Wang et al., 2019). To address the recruitment of AtEH/Pan1 and TPLATE in endocytic foci at the PM, we combined AtEH/Pan1 with TPLATE in their respective complemented mutant backgrounds. Similar to the combination of the core subunits, both AtEH1/Pan1-mRuby3 and AtEH2/Pan1-mRuby3 showed a very high degree of colocalization with TPLATE-GFP at the PM (Fig. 4D and E). To exclude the possibility that TPLATE and AtEH/Pan1 foci overlapped due to the high density of endocytic foci, we compared the colocalization percentage between TPLATE and AtEH1/Pan1 on images where the TPLATE channel was rotated 90 degrees. This rotation resulted in only around 10% of foci colocalizing (Fig. 5F), confirming that the observed high degree of colocalization is not random. These results also suggest a tight association between TPLATE and the AtEH/Pan1 proteins at the PM.

**Figure 5.**
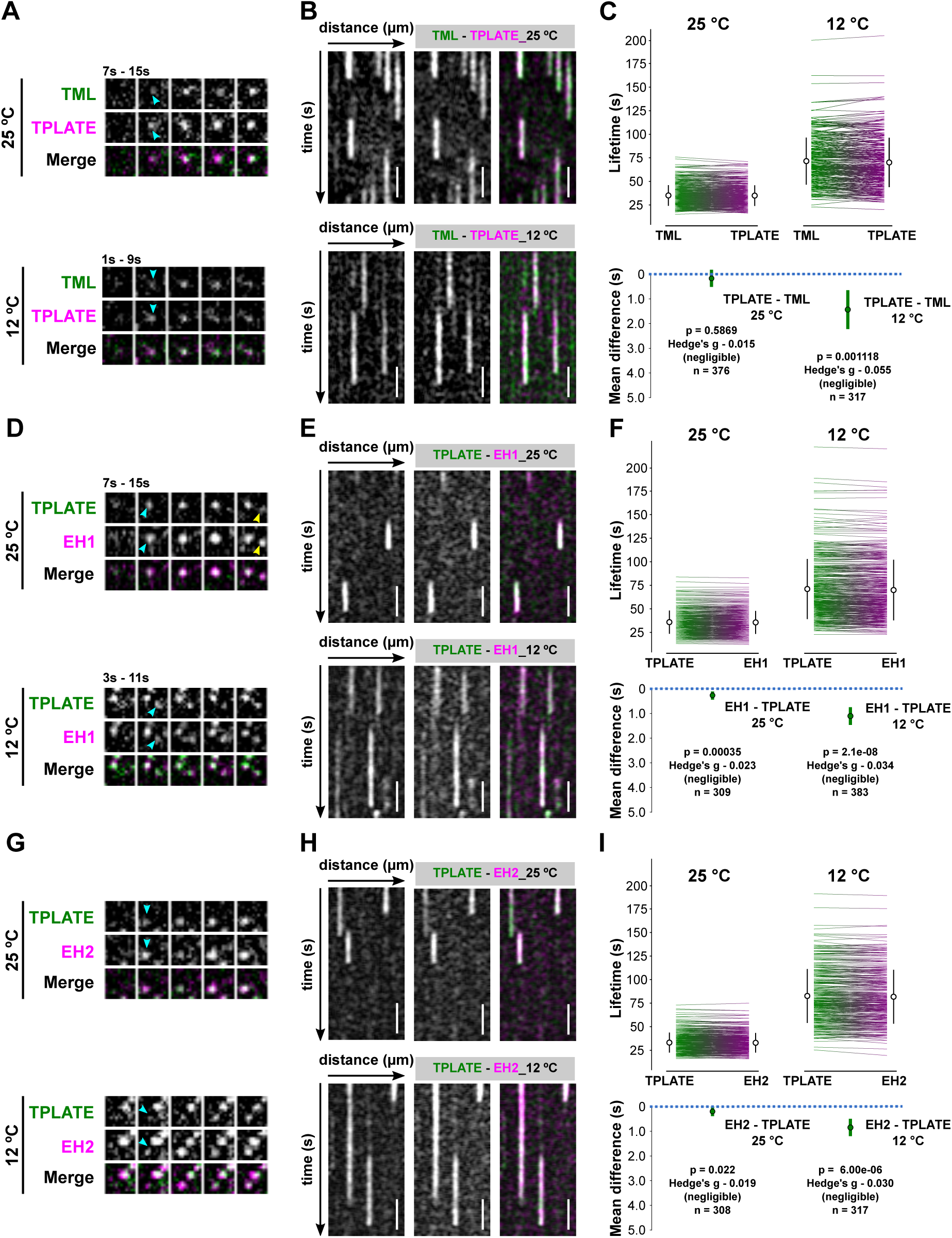
The AtEHs/Pan1 and the core TPC subunits are simultaneously recruited to the PM. (A, D and G) Representative time series of dual-color spinning disk movies (2s/f) showing the recruitment between TPLATE and TML, EH1/Pan1 and EH2/Pan1 at different temperatures. Arrows mark the appearance of TPLATE or TML, EH1/Pan1 and EH2/Pan1 on the PM. (B, E and H) Representative kymographs displaying the recruitment of TPLATE versus TML, EH1/Pan1 and EH2/ Pan1 at different temperatures. Scale bar, 25 µm. Time, 120s. (C, F and I) Paired comparison of the life-times of particular protein pairs at different temperatures. Each line on the sides of each plot represents an individual pair with the mean (white circle) ± SD (black line) for different combinations of constructs and temperatures. The p-value was calculated by the Wilcoxon Signed Rank test. The green circle in the bottom graph represents the paired mean difference with the bootstrap 95% confidential interval (green line). Hedges’ g value is a standardized effect size. n represents the number of events.

### AtEHs/Pan1 and the core TPC subunits are simultaneously recruited to the PM

To further investigate TPC assembly at the PM at the level of the different subunits, we compared the recruitment and disassociation behavior among pairs of TPC subunits at 25 and 12 °C. As a benchmark, we compared the behavior of two closely related TPC subunits, TPLATE and TML, which by homology to other adaptor complexes are presumed to be part of the core of TPC.

Time-lapse imaging and kymograph analyses showed that dual-labeled TML-GFP and TPLATE-TagRFP foci appear and disappear simultaneously at the PM when imaged at 25 °C (Figure 5A, 5B and S6). Lowering the temperature maintained this simultaneous appearing and disappearing behavior. However, there were very small changes in average lifetime, which were found to be negligible by their effect size (expressed as the Hedge’s g value). These small changes could be explained by there being more signal noise and differential bleaching effects of the fluorophores during the image acquisition at the lower temperature conditions (Figure 5A-5C and S6).

We next compared the PM appearance and disappearance behavior of AtEH/Pan1 and TPLATE. Similar to the recruitment behavior of TPLATE and TML foci, when imaged at 25 °C, both AtEH1/Pan1 and AtEH2/Pan1 foci simultaneously appeared and disappeared with TPLATE at the PM (Fig.D-E, G-H and S7). Lowering the temperature did not alter the simultaneous recruitment of TPLATE and AtEH/Pan1, and the respective Hedge’s value remained very low (Fig.D-I and S7). These results, together with our colocalization analysis, strongly suggest that AtEH/Pan1 subunits are not individually recruited to, or maintained, at the PM.

## Discussion

Clathrin-mediated endocytosis is a highly dynamic process. It is best understood in mammalian and yeast model systems while little is known in plants. Although very fast endocytic events were observed in neurons (Balaji and Ryan, 2007), in yeast or cultured animal cells, the whole endocytic process takes up to minutes and the physiological role and precise temporal dynamics of endocytic proteins are well defined (Lu et al., 2016; Ma et al., 2016; Pascolutti et al., 2019; Picco and Kaksonen, 2018; Taylor et al., 2011; Wrobel et al., 2019). In plant cells, the endocytic process is much more dynamic, with larger clathrin cages being formed than those in yeast in shorter time periods (Narasimhan et al., 2020), which brings about more technical difficulties in monitoring the temporal recruitment of the diverse endocytic players (Zhang et al., 2015). To better understand the dynamics of endocytosis in plant cells, new approaches are required, which should provide us with enhanced temporal resolution.

Here, we report that lowering the temperature using a microfluidics-based on-slide approach can be used as a rapid and efficient tool to slow down intracellular dynamic processes in plant cells similar to earlier reports demonstrating temperature effects on endocytosis and synaptic vesicle recycling in animal cells (Bui and Glavinovic, 2014; Pyott and Rosenmund, 2002). The effect on microtubule growth rate was immediate and the dampening effect on the dynamic of endocytosis, as measured by the dwell time at the PM of several markers, FM-uptake as well as measuring recruitment by FRAP correlated with the reduction in temperature. Lowering the temperature however had no significant effect on the density of endocytic foci, suggesting that cargo proteins could still be adequately recognized.

Our data also show that the temporal resolution of the differential recruitment of endocytic players could be enhanced at lower temperatures, which was most apparent between early and late arriving endocytic proteins. Lowering the temperature also enabled us to slightly enhance the temporal difference between two early arriving proteins, TPLATE and CLC2, indicating that our approach worked. Given the temperature effect on the dynamics of plant processes, awareness should be increased to perform live-cell imaging in plants under accurate temperature control. No doubt this will substantially increase the robustness and reproducibility of future image data collection.

The TPC is an evolutionary ancient adaptor complex, which was not retained in yeast or animal cells (Hirst et al., 2014). It is proposed to function as an adaptor complex during CME and has been shown to be recruited to the PM earlier than Dynamin-related proteins while only slightly earlier than clathrin(Gadeyne et al., 2014; Narasimhan et al., 2020). *Dictyostelium discoideum* contains a similar complex, which however lacks the AtEH/Pan1 proteins and in contrast to TPC, TSET is not essential (Gadeyne et al., 2014; Hirst et al., 2014). These AtEH/Pan1 proteins both share their male sterility phenotype with the other TPC subunits (Gadeyne et al., 2014; Van Damme et al., 2006; Wang et al., 2019), they are dynamically recruited to the PM (Gadeyne et al., 2014; Wang et al., 2019), associated with the other TPC subunits as well as AP-2, but not when TPC is forced into the cytoplasm using proteomic manipulations (Gadeyne et al., 2014) and functionally associated with several endocytic players during autophagy (Wang et al., 2019). Here we investigated how these AtEH/Pan1 proteins are recruited to the PM in relation to the other TPC subunits. Our live-cell analysis could not identify any deviation in recruitment dynamics between AtEH/Pan1 and TPLATE, mirroring the results observed between the other core subunits TPLATE and TML. The small differences reported could be attributed to technical issues, including differential bleaching effects of fluorophores. The close association, as well as the simultaneous recruitment between TPLATE and AtEH/Pan1 labeled endocytic foci at the PM, therefore allow us to hypothesize that TPC is recruited to the PM as an octameric complex in plants.

In plants, lifetimes of endocytic proteins are measured using time-lapse imaging of cells expressing fluorescent labeled proteins imaged via spinning disk microscopy (Bashline et al., 2013; Bashline et al., 2015; Dejonghe et al., 2016; Dejonghe et al., 2019; Gadeyne et al., 2014; Zhou et al., 2018) or total internal reflection fluorescence microscopy (TIRF) or variable angle epifluoresence microscopy (VAEM) (Ito et al., 2012; Johnson and Vert, 2017; Konopka et al., 2008; Konopka and Bednarek, 2008a, b; Narasimhan et al., 2020). Based on earlier work in the animal field (Aguet et al., 2013; Compeer et al., 2018; Jaqaman et al., 2008; Pascolutti et al., 2019; Wrobel et al., 2019), automated image analysis tools have recently been employed to analyze CME images unbiasedly and rapidly (Johnson and Vert, 2017; Narasimhan et al., 2020). Accurate detection of endocytic events requires high signal-to-noise ratio (SNR) images, a quality which is determined by multiple factors such as the susceptibility of the fluorophore to bleaching and the expression level of the fluorescent reporter and close-to-endogenous levels of fluorescent labeled endocytic proteins often failed to provide sufficient SNR for accurate lifetime measurements (Aguet et al., 2013). Here, we aimed at visualizing the recruitment dynamics of various TPC subunits at the PM. In order to avoid competition with endogenous subunits and to avoid over-expression effects such as induction of autophagy (Wang et al., 2019), we opted to work with double-labeled endocytic proteins in either single or double complemented *tpc* mutant background rather than use overexpression lines. Although our approach represents the optimal strategy from a biological perspective, our transgenic lines, especially those fused with red fluorescent proteins, and combined with extended imaging times under low temperature conditions failed to generate sufficient SNR images for automatic quantification. Actually, the only combination which yielded acceptable data when automatically quantified was our TPLATE-DRP1A combination, as the DRP1A is actually 35S-driven.

We therefore generated kymographs of our dual-color endocytic spinning disc movies and measured the dwell-times of the proteins manually. We aimed at measuring several hundred events per experiment and combined measurements of independent persons to achieve an unbiased assessment of the data. We were unable to identify any recruitment of AtEH/Pan1 proteins at the PM which was independent of another TPC subunit. Our findings therefore show that the TPLATE adaptor protein complex is likely recruited to the PM as an octameric unit, both at normal and at lowered temperature conditions. We can however not exclude the possibility that the differential recruitment between the individual TPC subunits are too dynamic to monitor under our limitating, one-second temporal resolution, conditions. However, due to the fact that we were able to detect differences in the CLC2 and TPLATE dynamics, which are of the order of a few seconds, we hypothesize that TPC is recruited as an octameric unit at the PM during CME. The continuous development of novel microscopy as well as labeling techniques will help to overcome our current limitations of working with low-expressing functional fusions and will allow us to understand how endocytosis in plants is executed at high spatiotemporal resolution.

## Materials and Methods

### Molecular cloning

To yield the expression construct for TWD40-1, entry clones of TWD40-1 without a stop codon in pDONR221(Gadeyne et al., 2014) were combined with pB7m34GW (Karimi et al., 2007), pDONRP4-P1R-Histone3p (Ingouff et al., 2017), and pDONRP2-P3R-mRuby3 (Wang et al., 2019) in a triple gateway LR reaction (Invitrogen) to generate pH3::TWD40-1-mRuby3. To yield a red fluorophore-tagged TPLATE, TPLATE without stop in pDONR207 (Van Damme et al., 2006) was combined with the Lat52 promotor in pDONRP4P1R (Van Damme et al., 2006), with TagRFP in pDONRP2P3R (Mylle et al., 2013) and with pH7m34GW (Karimi et al., 2007) in a triple gateway LR reaction (Invitrogen) to generate pLat52::TPLATE-TagRFP.

### Plant Material

The *Arabidopsis* lines expressing EB1a-GFP (Van Damme et al., 2004), LAT52p::TPLATE-GFP (Gadeyne et al., 2014; Van Damme et al., 2006), TMLp::TML-GFP (Gadeyne et al., 2014), 35Sp::DRP1a-mRFP (Mravec et al., 2011) and 35Sp::CLC2(At2g40060)-mKO (Ito et al., 2012) were previously described. The dual-color line expressing TPLATE-GFP/CLC2-tagRFP in *tplate* homozygous background was reported previously (Gadeyne et al., 2014). The dual-marker lines expressing TPLATE-GFP/AtEH1/Pan1-mRuby3 or TPLATE-GFP/AtEH2/Pan1-mRuby3 in double homozygous mutant (*tplate ateh1/pan1* or *tplate ateh2/pan1*) background were reported before (Wang et al., 2019).

To generate TWD40-1-mRuby3 complemented lines, heterozygous mutants of *twd40-1-1* (Gadeyne et al., 2014) were transformed with pH3.3::TWD40-1-mRuby3 by floral dip. The T1 plants were selected for the complementation constructs on 1/2 MS plate supplemented with 10mg/L Basta. Resistant plants were genotyped to identify those with a *twd40-1-1* heterozygous mutant background. T2 plants expressing TWD40-1-mRuby3 were tested by genotyping PCR to identify homozygous lines for the *twd40-1-1* insertion mutations. Genotyping PCR was performed on genomic DNA isolated from rosette leaves. Genotyping primers for *twd40-1-1* are described before (Gadeyne et al., 2014).

For backcross experiments, the complemented lines of TWD40-1-mRuby3 as well as the heterozygous mutant plants of *twd40-1-1* were used as male to cross with Col-0 as female. The transfer of the T-DNA, causal to the functionality of the complementing fusion, was analyzed by genotyping PCR on F1 plants.

To generate dual-marker lines of TPLATE core subunits, a complemented *tplate* lines expressing pLat52::TPLATE-tagRFP was generated by dipping tplate(+/-) heterozygous plants with pLat52::TPLATE-TagRFP and selecting for the transformants. Homozygous *tplate(-/-)* plants were identified in the next generations by genotyping PCR.

Homozygous *tplate* mutant plants carrying pLat52::TPLATE-TagRFP were crossed with the complemented *tml-1* line expressing TMLp::TML-GFP (Gadeyne et al., 2014). The complemented *twd40-1-1* lines expressing TWD40-1-mRuby3 were crossed with the complemented *tplate* line Lat52p::TPLATE-GFP, respectively. F2 plants in double homozygous mutant background (*tml-1/tplate* or *tplate/twd40-1-1* were identified by genotyping PCR. For the dual-color TPLATE-GFP/DRP1a-mRFP lines, the complemented *tplate* lines expressing Lat52p::TPLATE-GFP were crossed with 35Sp::DRP1a-mRFP expressing lines and F2 plants in *tplate* homozygous background were identified by genotyping PCR. Genotyping primers for *tplate, tml-1 and twd40-1-1* are described before (Gadeyne et al., 2014).

### Temperature modulation using the CherryTemp System

The CherryTemp system (Cherrybiotech), which enables ultra-fast temperature shifts between 5 °C and 45°C was used to modulate and maintain the temperature during the spinning disk imaging (https://www.cherrybiotech.com/heater-cooler-for-microscopy). Prior to imaging, a single etiolated seedling was laid between two coverslips with 1/2 strength MS liquid medium and incubated with the CherryTemp Heater Cooler device for five minutes prior to imaging to stabilize the temperature of the seedling (https://www.cherrybiotech.com/heater-cooler-for-microscopy/temperature-control-for-plant-microscopy).

### FM4-64 Uptake

Prior to imaging, whole 6-day-old Col_0 seedlings were incubated with 4 μM FM4-64 (Invitrogen) solution in 1/2 strength MS liquid medium between 2 coverslips at 25 °C or 12 °C for 30 minutes.

### Live-cell Imaging

A Nikon Ti microscope with the Perfect Focus System (PFSIII) for Z-drift compensation, equipped with an Ultraview spinning-disk system (PerkinElmer) and two 512×512 Hamamatsu ImagEM C9100-13 EMccd cameras was used to image endocytic dynamics. Images of hypocotyl epidermal cells of 4-day old etiolated seedlings expressing single or dual-color fluorophore labeled proteins were acquired with a 100x oil immersion objective (Plan Apo, NA = 1.45). During imaging, the CherryTemp system was used to maintain the temperature of the samples constant.

Seedlings expressing GFP fused proteins were imaged with 488nm excitation light and an emission window between 500 nm and 530 nm in single camera mode, or 500 to 550 nm in dual camera mode. Seedlings expressing mKO, mRFP and tagRFP labled proteins were imaged with 561 nm excitation light and an emission window between 570nm and 625nm in single camera mode or 580 to 630 nm in dual camera mode. Single-marker line movies were acquired with an exposure time of 500 ms/frame. Movies of 2 minutes in total were made. Dual-color lines were acquired either sequentially (one camera mode) or simultaneously (two camera mode) with an exposure time of 500 ms/frame. Single camera mode was used for density, colocalization (TML-TPLATE, TPLATE-DRP1A) and life-time (TML-DR1AP,TML-TPLATE 25°C) measurements. Dual camera mode was used for colocalization (TPLATE-AtEH1/Pan1, TPLATE-AtEH2/Pan1, TPLATE-TWD40-1, TPLATE-TWD40-2) and lifetime (TPLATE-AtEH1/Pan1, TPLATE-AtEH2/Pan1 and TML-TPLATE 12°C) measurements. For photo-bleaching experiments, seedlings were exposed to 100% power of laser for around 2s during the imaging, using the Photokinesis unit of the PE spinning disk system.

Root meristematic epidermal cells of 6-day-old seedlings were acquired with a Zeiss 710 inverted confocal microscope with the ZEN 2009 software package and equipped with a C-Apochromat 40x water Korr M27objective (NA 1.2). FM4-64 was visualized using 561 nm laser excitation and a 650–750 nm detection window.

### Quantification of measurements

Dynamics of EB1a-GFP were analyzed using ImageJ equipped with the Trackmate (v4.0.1) plugin(Tinevez et al., 2017). We used the LoG detector with an estimated blob diameter of 10 pixels, a threshold value of 1, using the median filter and sub-pixel localization. An auto initial thresholding, followed by a Linear motion LAP tracker with as parameters a 5 pixel initial search radius, a 5 pixel search radius and a 2 frames maximum frame gap correction, were applied. Tracks with a duration of less than 10 frames were excluded and the obtained median speed per track was converted to µm/min using the pixel size values. Outliers for the median speed were defined by the iterative Median Absolute Deviation Method (MAD) (Leys et al., 2013) and their values were excluded. Three movies coming from two different seedlings were analyzed.

The density of labeled endocytic markers was measured in Matlab 2017b using the detection and tracking parts of the of the cmeAnalysis package and further processed as described in (Johnson and Vert, 2017; Narasimhan et al., 2020). Density calculations were obtained from all the tracks within the region of interest (ROI) over certain frames of the movies, which used to produce an average density. The ROI was selected on the middle of the image. The middle frame was used as a reference and the temporal range is based on the middle frame. We used the pixel size and the area to convert this to spots per µm^2^. For each of the analyzed sample set, eight cells from eight different seedlings were analyzed.

Objects based co-localization was performed using the ImageJ plugin Distance Analysis (DiAna) (Gilles et al., 2017). Prior to analyzing with DiAna, images were processed with ImageJ. Each channel was processed with a Walking Average of 4 and then merged (also rotated if required). Region of interest were selected based on that they excluded the border of the cells and still represented a good number of objects. Z-projection images were generated of five slices with average intensity. Each channel of Z-projected images was processed using Morphological filters from the MorphoLibJ plugin (Legland et al., 2016), using the parameters white top-hat, disk element and a 2 pixels radius. A first step in the DiAna plugin is to segment the objects for each channel, which is done by selecting the 3D Spot segmentation tool of the DiAna-Segment plugin. If requested, adapt the calibration by changing the pixel size to 1.00001 for all dimensions. Both the noise and seed threshold value were obtained by averaging the maximum intensity of three regions covering only background signal. The spot was defined with a minimum value of 4 and maximum value of 36 pixels. The option to exclude objects on XY edges was activated. Default values were used for the other parameters. Results for number of total objects (Tot) or touching objects (Tou) in image A/B obtained from Diana were recorded. The colocalization ratio of objects was calculated as follows:

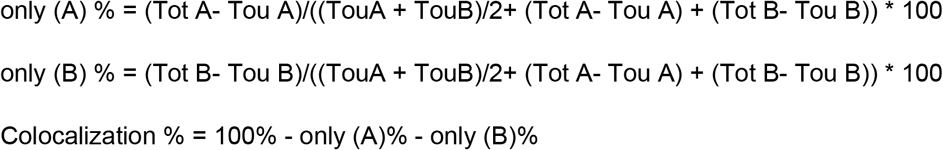

As a control, one of the channels was rotated 90°C (no interpolation) and analyzed similarly as described above. For each of the analyzed sample set, a minimum of six cells coming from three different seedlings were analyzed.

Lifetimes of individual endocytic events were measured from kymographs generated by the Volocity software package (PerkinElmer). To measure paired lifetimes of dual-color kymographs, individual events showing good SNR (signal-noise ratio) in both channels were marked. Following the measurement of the lifetimes of the marker in the red channel, the lifetime of the marker in the green channel was analyzed. Data was further analyzed in Excel by checking the start position of X from each line of the kymograph to avoid mistakes in pairing the red and green channel values. Calculation to time (ns) of each line was done. Outliers for the life-time differences were defined by the iterative MAD (Leys et al., 2013) and their values were excluded. For each of the analyzed sample sets, minimum 9 movies spread among minimum 3 seedlings were analyzed. Except for TPLATE-GFP/CLC2-tagRFP for which 7 movies over 3 seedlings (12°C) and 5 movies over 2 seedlings (25°C) were analyzed. To generate unbiased data, paired lifetimes of endocytic events labeled by dual-color fluorophores were measured by independent persons.

### Statistical analysis

The results were analyzed with the estimation method to calculate mean, mean differences, confidence intervals, and Hedges’ g (Claridge-Chang and Assam, 2016; Ho et al., 2019). 95% confidence intervals for the mean differences were calculated using bootstrap methods (re-sampled 5000 times, bias-corrected, and accelerated). Effect size was measured using Hedges’ g and accordingly to the standard practise is referred as ‘negligible’ (g < 0.2), ‘small’ (0.2 < g < 0.5), ‘medium’ (0.5 < g < 0.8) or ‘large’ (g > 0.8) (Cumming, 2012). Hedges’ g is a quantitative measurement for the difference between mean indicating how much two groups differ from each other, if Hedges’ g equals 1, the two groups differ by 1 standard deviation. R version 3.6.2 and Rstudio 1.2.5001 were used to calculate the Wilcoxon Signed Rank test (paired samples), the Mann-Whitney U-test and the Welch two sample T-test (Team, 2019). All plots and figures were generated with Rstudio 1.2.5001 and Inkscape (version 0.92, https://inkscape.org/).

## Supplemental Material

Figure S1. The CherryTemp system can be effectively used to quickly alter intracellular dynamics in plant cells.

Figure S2. Lowering the temperature increases the lifetime of endocytic proteins at the plasma membrane.

Figure S3. Lowering the temperature decreases the recruitment dynamics of the endocytic proteins.

Figure S4. Lowering the temperature enhances the temporal resolution of recruitment.

Figure S5. TWD40-1 localizes to the PM as endocytic foci.

Figure S6. TPLATE and TML are recruited to the PM simultaneously.

Figure S7. AtEHs and TPLATE are recruited to the PM simultaneously.

## Acknowledgements

Research in the Van Damme lab is funded by the European Research Council T-Rex project number 682436 and by the Chinese Scholarship Council (CSC) (scholarship 201508440249 to J.W.). A.J. is supported by funding from the Austrian Science Fund (FWF): I3630B25 to J.F.

## Declaration of Interests

The authors declare no competing interests.

